# Pupil and Neural Dynamics Reveal Belief-Dependent Decision Making Under Ambiguity

**DOI:** 10.64898/2026.04.21.719818

**Authors:** Yinuo Qin, Nuttida Rungratsameetaweemana, Nina Lauharatanahirun, Paul Sajda

## Abstract

Humans make thousands of decisions under ambiguity every day, where most times the probabilities of getting beneficial outcomes are not fully known. Although ambiguity aversion is well characterized at the aggregate level, it remains unclear how individuals internally represent ambiguous outcomes and how these representations shape behavior and physiology. To address these questions, we investigate value-based decision-making under risk and ambiguity using a multimodal approach that combines behavioral, pupillometric, and electroencephalographic (EEG) data. We show that individuals adopt distinct decision strategies that reflect different internal beliefs about unknown outcomes. These belief differences are expressed not only in choice but also in pupil-linked arousal during decision formation. Crucially, physiological differences across strategies persist under an objective valuation rule for ambiguous options but disappear when valuation is computed using each individual’s subjective beliefs, indicating that arousal tracks subjective belief rather than objective task structure. Together, our findings show that ambiguity aversion is not a uniform bias but reflects heterogeneous internal belief models, and that multimodal physiology provides a window into the mechanisms through which these beliefs guide value-based choice.

## 1 Introduction

Humans make decisions in situations where outcomes are unknown, and the probabilities of benefit and harm are incomplete or unknown. Such decisions shape behavior across many domains, including financial investment, medical decision making, social interaction, and collaboration with artificial intelligence systems. A key distinction is between *risk*, where outcome probabilities are known, and *ambiguity*, where outcome probabilities cannot be precisely determined (15; 37; 36). For example, risk is betting on a fair coin toss, whereas ambiguity is betting on a coin drawn from a set of coins with different and unknown biases. Ambiguity poses a distinct cognitive challenge and often elicits behavioral responses that differ from those observed under risk (37; 3; 64). Understanding how individuals interpret and respond to ambiguity, and the underlying computational and neurophysiological mechanisms that give rise to these responses, is therefore central to explaining human decision-making in naturalistic settings.

A central distinction in decision theory separates decisions under risk from decisions under ambiguity. Under risk, outcome probabilities are known; under ambiguity, they are partially or entirely unknown. The behavioral consequences of this distinction were first formalized by the Ellsberg paradox, which showed that people systematically prefer risky options with known probabilities over ambiguous options with unknown probabilities, even when the expected values are equivalent (18). Subsequent work across economics, psychology, and neuroscience has consistently demonstrated ambiguity aversion, often interpreted as a general bias against unknown probabilities (14; 3; 11). However, much of this literature focuses primarily on choice behavior, characterizing how often individuals avoid ambiguity, while leaving open questions about how ambiguity is internally represented and processed during decision formation.

Despite robust population-level evidence for ambiguity aversion, aggregate choice patterns can obscure substantial inter-individual differences. People vary widely in risk tolerance, learning strategies, and how they form beliefs about unknown outcomes (19). Yet ambiguity aversion is often treated as a single bias or a fixed trait that applies uniformly across individuals (1; 14; 33; 31). This perspective overlooks the possibility that different people may hold qualitatively different internal representations of ambiguity, even when they face the same objective risk. Moreover, population-level analyses rarely link behavioral variability to corresponding physiological or neural responses that could reveal how ambiguity is processed internally. As a result, it remains unclear what individuals actually believe when probabilities are unknown.

Physiological measures offer a powerful window into the internal processes that underlie decision-making under risk. Pupil size provides a sensitive index of arousal and valuation, particularly in value-based decision tasks where it tracks risk, surprise, and subjective value (45; 22; 56; 62; 23). Electrophysiological signal complements this by revealing the temporal dynamics of neural processes related to control, attention, and valuation during decision formation (27; 35; 44; 50; 40). Prior work has linked both pupillary and EEG signals to uncertainty, cognitive conflict, effort, and attentional engagement (26; 9; 61). However, these physiological signatures are rarely combined with explicit inference of individuals’ internal beliefs about ambiguous outcomes. As a result, it remains unclear whether physiological signals reflect objective properties of risk or the subjective beliefs individuals hold when probabilities are unknown.

Behavioral choices alone provide an incomplete picture of how decisions under ambiguity are formed. Similar choices can arise from different internal beliefs or computational strategies, making it difficult to infer how risk is represented from behavior alone (14; 19). Trial-level physiological measures offer a complementary approach by capturing internal states as decisions unfold. However, it remains unclear whether and how these physiological dynamics reflect the latent belief models that individuals use to interpret ambiguous outcomes, and whether they can explain stable individual differences in ambiguity processing beyond what is observable from aggregate choice patterns.

To address these questions, we developed a value-based decision-making task with controlled levels of ambiguity and simultaneously multi-modal physiological recordings. Participants chose between a sure option and a lottery with partially unknown outcome probabilities while pupil size and EEG were measured throughout the decision process. Using behavior under non-ambiguous conditions, we characterized participants into three decision styles (*ideal, aggressive*, and *conservative*) based on how their choices aligned with expected-value optimality. We then inferred subjective belief parameters that captured how individuals internally interpreted the ambiguous probability mass and examined how these beliefs related to choice behavior, arousal dynamics, and neural activity. This multimodal framework allowed us to study ambiguity processing at the level of individual belief models rather than aggregate choice patterns.

We identify distinct decision styles under ambiguity that reflect stable internal belief models rather than random variability or uniform bias. Our results indicate that behavioral asymmetries across individuals are explained by differences in subjective beliefs about ambiguous outcomes: *aggressive* participants exhibit optimistic valuation, *conservative* participants exhibit pessimistic valuation, and *ideal* participants occupy an intermediate regime. Crucially, physiological signals validate these belief models. Pupil-linked arousal tracks subjective valuation rather than objective expected value, and neural dynamics reveal belief-specific computational regimes involving control, excitability, and engagement. Together, these findings provide a belief-level and computational account of how ambiguity is represented, evaluated, and acted upon by individuals, linking internal beliefs to behavior and physiology in value-based decision making. Notably, individual decision strategies expressed in this non-collaborative task are also associated with leadership tendencies observed in prior team-based interactions, suggesting broader relevance beyond isolated choice behavior.

## 2 Results

### 2.1 Task Structure and Behavioral Stratification

Participants first completed a three-person collaborative spacecraft control task requiring real-time coordination across distinct degrees of freedom under partial and complementary visual information (48). They then individually performed a value-based decision-making task (Lottery Choice Task, LCT; Fig. 1a) designed to dissociate decision formation under *risk* and *ambiguity*.

**Fig. 1:**
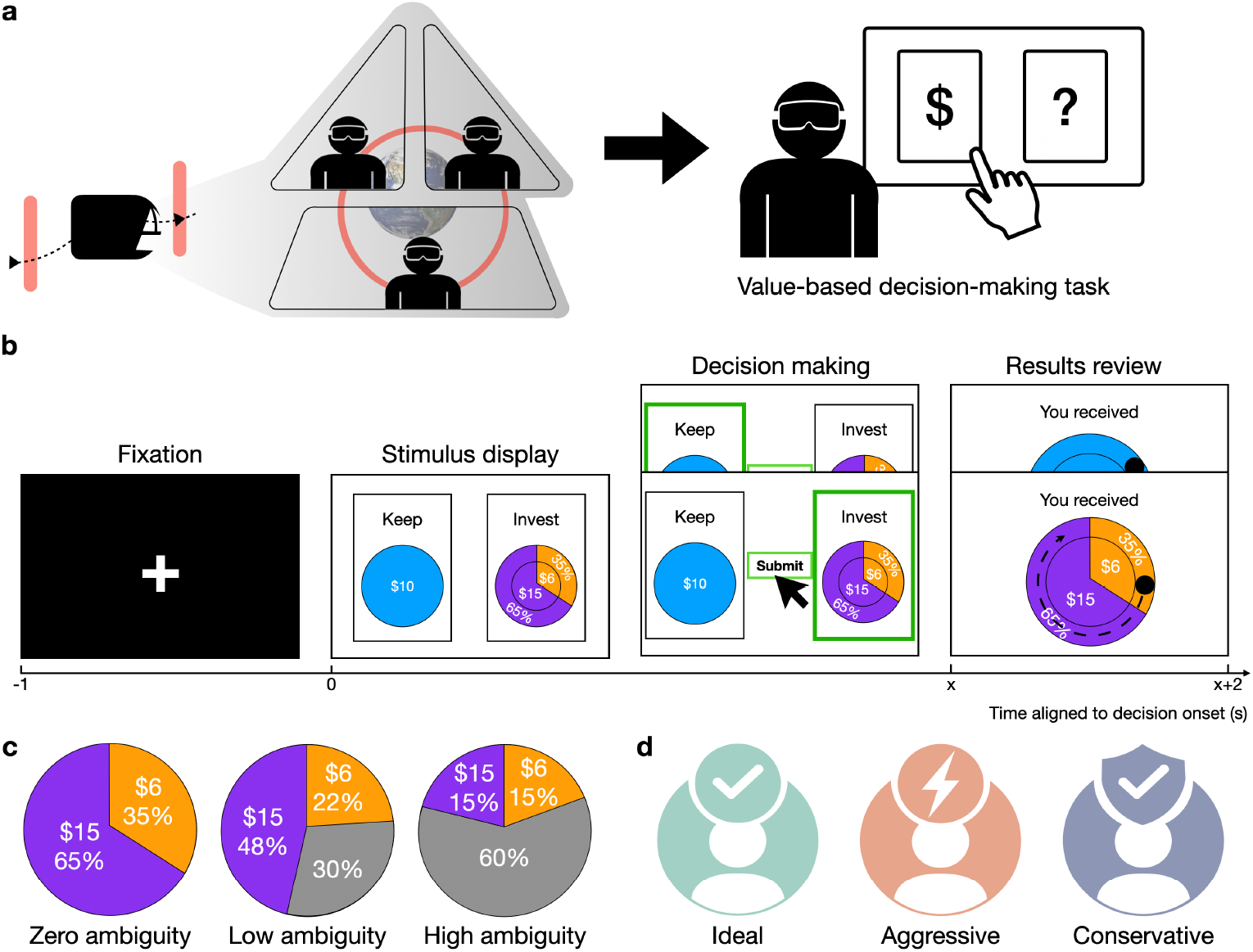
Experimental setup. **a**, Three participants first perform a collaborative space craft control task. In this task, three participants with different perspectives control a single degree of freedom of the spacecraft movement toward earth, coordinate through communication while avoiding obstacles. Afterward, each participant completed the Lottery Choice Task (LCT) individually. **b**, The temporal structure of a single trial of the LCT. The horizontal axis denotes time in seconds, and analyses were time-locked to stimulus onset. Each trial started with a 1 s fixation. Then, the stimulus was presented, and participants had up to 5 s to make a *keep* or *invest* decision (*x* ≤5). Outcome feedback was then revealed for 2 s following the decision, after which the next trial began with a fixation period. The values and risk probabilities shown in panels **b** and **c** are illustrative examples. **c**, Trials varied in ambiguity level. Under zero ambiguity, risk probability is fully known to participants. Under low (30%) or high (60%) ambiguity, partial risk is unknown when making decisions. **d**, Based on participants investment decisions across trials, they were categorized as *Ideal, Aggressive*, or *Conservative*.

On each trial, participants chose whether to *keep* a sure endowment or *invest* it into a lottery whose outcomes consisted of an explicit gain and loss (Fig. 1b). In zero ambiguity (**risk** only) trials, the gain, loss, and their associated probabilities were fully specified, allowing the expected value (EV) of the investment option to be computed from complete information. In **ambiguity** trials, the payoff magnitudes remained explicit, but the probability of receiving the gain or loss was only partially specified: the task masked a portion of the probability mass, leaving participants with an interval rather than a single known probability (Fig. 1c). Ambiguity differs fundamentally from risk at both the physiological level (34; 6) and the behavioral level (37; 4).

Throughout the paper, we treat participants’ investment behavior under zero ambiguity trials as a behavioral proxy for decision strategy. Under this framing, different decision strategies reflect distinct decision policies governing valuation and action selection. To quantify inter-individual variation, we classified participants into three investor types based on how their choices aligned with the expected-value–optimal action across risk trials (Fig. 1d; see *Materials and Methods, Section 4.3*). This procedure yielded three exclusive groups: *ideal, aggressive*, and *conservative* investors. This stratification operationalizes individual decision policy using stable differences in investment behavior under known risk, enabling subsequent analyses of how these strategies generalize to ambiguity and shape physiological responses.

### 2.2 Behavioral and Physiological Response to Different Ambiguity

We examined how ambiguity modulated investment behavior across decision strategies (Fig. 2a). Consistent with classical ambiguity aversion as formalized in the Ellsberg paradox (18; 14), increasing ambiguity was associated with a systematic reduction in the probability of choosing to invest. This effect was pronounced in both *ideal* and *aggressive* investors: the majority of participants in these two groups showed a steep decline in investment rates as ambiguity increased from 0% to 60% (Fig. 2b, *χ*^2^ tests for ambiguity effects: 78.12% of *ideal* participants and 84.38% of *aggressive* participants exhibited a significant effect, *P <* 0.05). In contrast, *conservative* investors showed substantially weaker sensitivity to ambiguity. Their investment probability was already low at 0% ambiguity and decreased only modestly as ambiguity increased (46.88%). As a result, under high ambiguity, investment behavior converged across groups, whereas under low ambiguity, *ideal* and *aggressive* investors remained substantially more willing to invest than *conservative* investors. These results indicate that ambiguity exerts strong and graded control over value-based decisions, but that this control is selectively attenuated in individuals with conservative decision strategies.

**Fig. 2:**
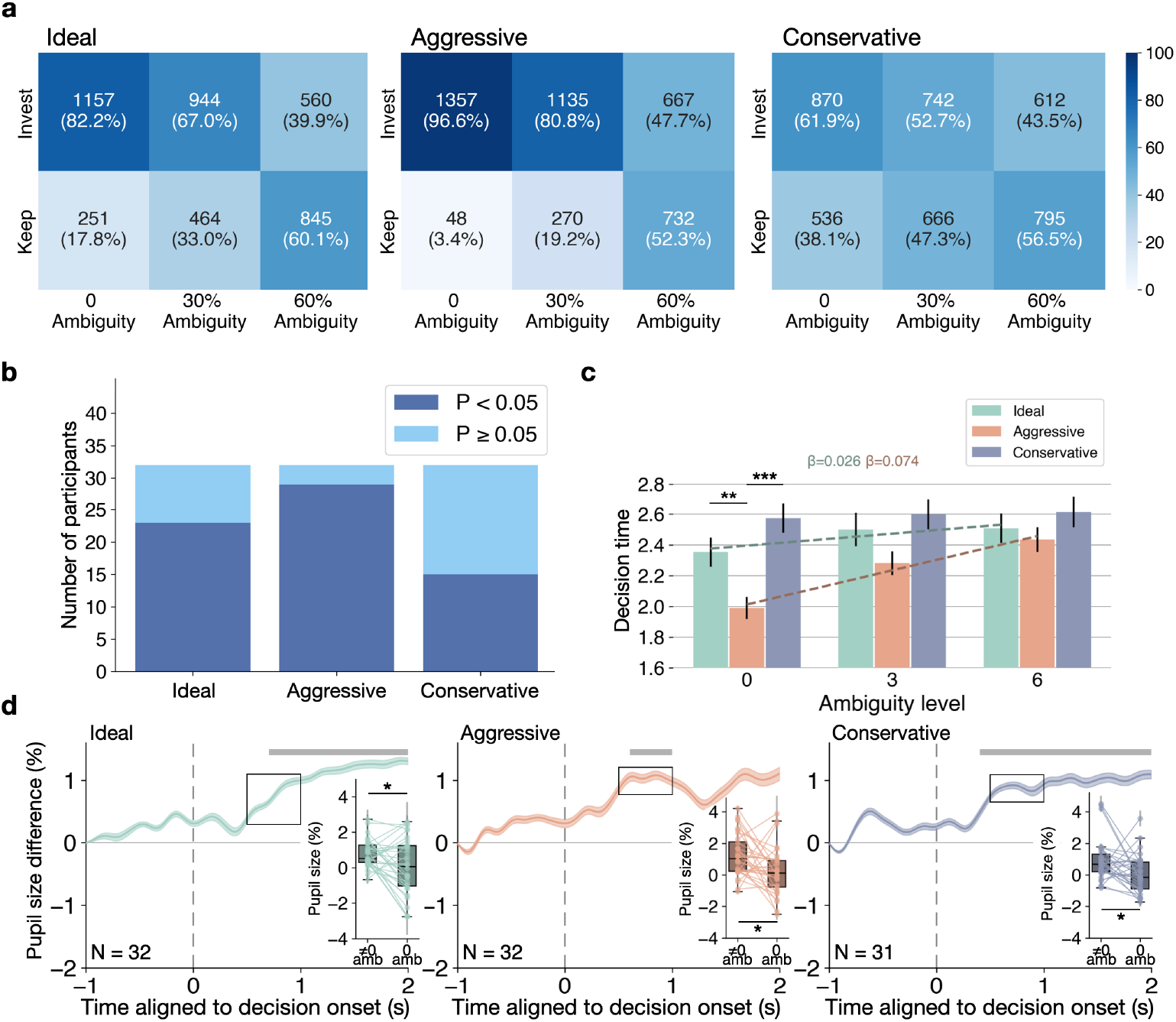
Behavioral and pupil signatures across decision strategies under ambiguity. **a**, Investment behavior across ambiguity levels (0%, 30%, 60%) for *ideal, aggressive*, and *conservative* investors. Heatmaps show the number and percentage of invest and keep decisions at each ambiguity level (*N* = 32 per group in panels **a**–**c**; total *N* = 96). **b**, Participant-level significance breakdown across strategy groups. Stacked bar plot showing the number of participants in each strategy group whose individual test result is statistically significant (*P <* 0.05, dark blue) versus non-significant (*P* ≥0.05, light blue) using *χ*^2^ test. **c**, Decision time as a function of ambiguity level and investor type. Bars indicate mean *±* standard error of the mean (SEM); dashed lines show fitted regression trends. Horizontal bars above the plots indicate significant group differences assessed using one-way ANOVA with posthoc *t*-tests and Bonferroni correction (0 Ambiguity: *F* (2, 93) = 12.71, *P <* 0.0001; *ideal* vs. *aggressive*: *T* (31) = 3.24, *P* = 0.0019, *aggressive* vs. *conservative*: *T* (31) = −5.38, *P <* 0.0001. 30% Ambiguity: *F* (2, 93) = 4.49, *P* = 0.0138; *aggressive* vs. *conservative*: *T* (31) = −3.08, *P* = 0.0031). **d**, Difference in pupil size between ambiguous and non-ambiguous trials over time for each investor type. Time 0 indicates decision start. Color-shaded regions denote SEM; gray bar on top indicate time intervals with significant effects evaluated using Wilcoxon signed-rank tests on sliding windows with false discovery rate (FDR) correction (*P <* 0.05). *N* denotes the number of participants per strategy group. Data in the black box are shown in box plots. The box plot with paired dots show each participant’s mean pupil size change from baseline (%) measured in 0.5-1 s after decision onset under non-zero ambiguity and zero ambiguity conditions. Each dot represents one participant’s averaged pupil response, and error bars indicate the within-participant SEM. Lines connect paired observations across conditions. Statistical significance was assessed using a paired *t*-test within each strategy group with Bonferroni correction (*ideal* : *T* (31) = 2.62, *P* = 0.0400; *aggressive*: *T* (31) = 2.74, *P* = 0.0304; *conservative*: *T* (30) = 2.63, *P* = 0.0401). Asterisks indicate significance as (\**P <* 0.05, * * *P <* 0.01, * * \**P <* 0.001).

Ambiguity also shaped the temporal dynamics of decision making (Fig. 2c). Across *ideal* and *aggressive* investors, decision time increased systematically with higher ambiguity (mixed effect model, *ideal β* = 0.03, *P <* 0.001; *aggressive β* = 0.07, *P <* 0.0001), indicating that resolving partially unknown probabilities required additional deliberation. In contrast, *conservative* investors exhibited consistently long decision times across all ambiguity levels, with no significant modulation by ambiguity (*β* = 0.01, *P* = 0.2745). As a result, group differences in decision time were stable as ambiguity increased. These findings suggest that while ambiguity imposes an additional temporal cost on the decision process for *ideal* and *aggressive* investors, *conservative* investors operate in a uniformly slow decision regime that is insensitive to changes in ambiguity.

Ambiguity further modulated pupil-indexed physiological arousal. For both *ideal* and *conservative* investors, ambiguous trials elicited significantly larger pupil size changes than non-ambiguous trials, indicating heightened arousal under conditions of partial ambiguity (Fig. 2d). This pattern is consistent with prior work linking pupil dilation to affective and arousal responses during decision making under ambiguity (45; 22; 56; 30). In contrast, *aggressive* investors showed no significant pupil modulation by ambiguity, with pupil responses remaining near baseline across the full decision window. This reduced physiological sensitivity align with shorter decision time (Fig. 2c) indicates that *aggressive* participants’ arousal level are less modulated by ambiguity. Together, these results suggest that ambiguity engages arousal systems in *ideal* and *conservative* investors, whereas *aggressive* investors exhibit a blunted arousal response to ambiguity that aligns with their stable, low-risk decision policy.

## 2.3 Physiological Response Differences for Different Investment Decisions

After establishing how ambiguity modulates behavior and pupil-linked arousal across decision strategies, we next examined how physiological responses differ between *keep*and *invest* decisions. Because participants’ decision strategies were defined using non-ambiguous trials, subsequent physiological comparisons were restricted to ambiguous trials to avoid circularity. This allowed us to isolate decision-specific physiological signatures under ambiguity, independent of the criteria used to characterize individual decision strategies.

### 2.3.1 Anticipatory and Choice-Dependent Pupil Arousal Signatures

Pupil size dynamics revealed robust decision-dependent arousal signatures that were consistent across decision strategies. In all three groups, pupil responses diverged systematically between *keep*and *invest* choices following decision onset, with *keep* decisions exhibiting larger pupil size changes than *invest* decisions throughout the subsequent two-second decision window (Fig. 3a). This time-locked separation indicates that pupil-linked arousal tracks evolving decision commitment rather than reflecting a static baseline difference. Such decision-dependent pupil modulation is consistent with prior work showing that pupil dilation reflects subjective belief states and uncertainty during value-based choices (45), and more broadly aligns with evidence linking task-evoked pupil dilation to phasic locus coeruleus–norepinephrine (LC–NE) activity during the decision process and behavioral engagement (58).

**Fig. 3:**
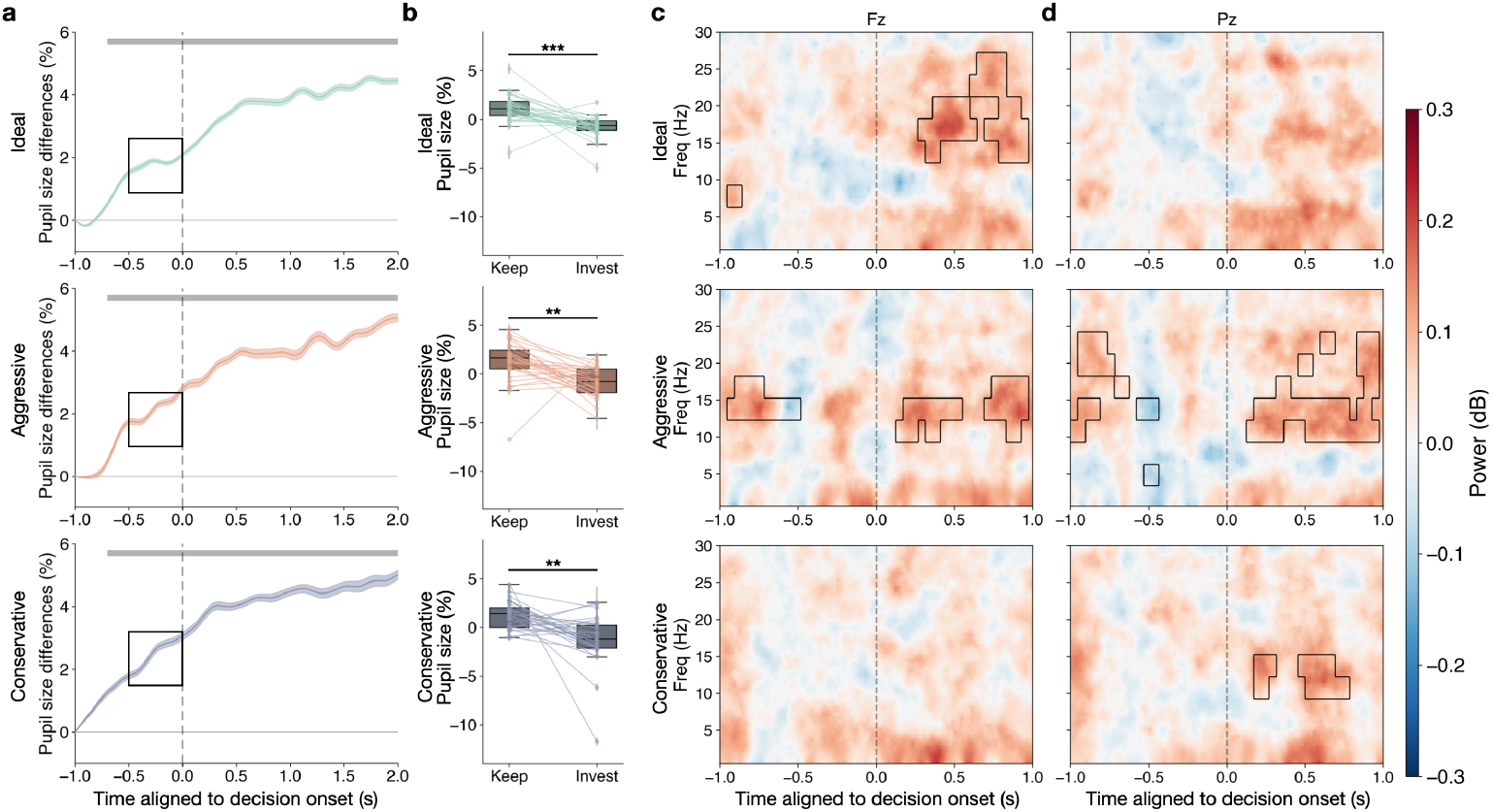
Decision-dependent pupil and EEG signatures across decision strategies. **a**, Time-resolved pupil size difference (*keep* minus *invest*) under ambiguous conditions for *ideal, aggressive*, and *conservative* investor groups. Shaded bands denote SEM across participants. Grey bar on top indicates time windows showing significant differences (*P <* 0.05), assessed using Wilcoxon signed-rank tests on sliding windows with false discovery rate (FDR) correction. **b**, Participant-level paired comparison of mean pupil size change from baseline (%) computed within the pre-decision interval highlighted by the black box in the left panels (0.5 s before decision onset). Each dot represents one participant (error bars: within-participant SEM), and lines connect paired observations. Group-level significance was assessed using paired *t*-tests with Bonferroni correction (*ideal* : *T* (30) = 4.82, *P* = 0.0001; *aggressive*: *T* (30) = 3.92, *P* = 0.0014; *conservative*: *T* (30) = 3.96, *P* = 0.0013). Asterisks denote significance (** *P <* 0.01, *\*\*\* P <* 0.001) and *N* = 31 for each strategy group in **a** and **b. c, d**, Time–frequency representations of EEG power differences (*keep* - *invest*) at frontal (Fz, **c**) and parietal (Pz, **d**) electrodes for each strategy group. Black outlines indicate time–frequency regions with significant differences (*P <* 0.05), evaluated using Wilcoxon signed-rank tests on sliding windows with FDR correction (*N* = 25 for *ideal* and *aggressive*; *N* = 29 for *aggressive*). Together, pupil and EEG results reveal distinct decision-dependent arousal and neural dynamics that vary systematically across *ideal, aggressive*, and *conservative* decision strategies.

Interestingly, pupil size differences between upcoming *keep*and *invest* choices were already present before the decision stimulus appeared. These differences indicate that arousal states predictive of choice emerged prior to the decision-making period (Fig. 3b). This anticipatory separation suggests that trial-to-trial variability in pupil-linked arousal reflects more than reactive processing of decision evidence (56; 17). These pre-stimulus signatures align with evidence that pupil dilation can track endogenous belief states and predict upcoming decisions, consistent with models linking pupil dynamics to phasic locus coeruleus–norepinephrine (LC–NE) activity that regulates attentional gain and behavioral engagement (45; 12; 58). Together, the presence of *keep*–*invest* pupil differences before stimulus onset supports an interpretation in which a preparatory neuromodulatory state contributes to shaping decision outcomes, rather than arising solely as a consequence of deliberation.

#### 2.3.2 Time–Frequency EEG Signatures of Strategy-Dependent Decision Process

EEG time–frequency analyses revealed reliable choice-dependent neural dynamics under ambiguity, with patterns that varied across decision strategies. We quantified spectrotemporal power differences between *keep*and *invest* decisions (keep – invest) at frontal (Fz, Fig. 3c) and parietal (Pz, Fig. 3d) electrodes, and identified significant time–frequency windows using sliding-window Wilcoxon signed-rank tests with false discovery rate (FDR) correction.

*Ideal* investors exhibited the most selective and temporally focused effects, dominated by post-decision frontal modulation. At the frontal channel, *ideal* investors showed a large, significant cluster spanning mid-to-high frequencies from about 0.3 s up to nearly 1.0 s after decision onset (Fig. 3c). This cluster reflected stronger beta-range activity during *keep* relative to *invest*, suggesting enhanced recruitment of frontal decision-related oscillatory dynamics during the post-choice interval (5; 13; 24). In addition, a smaller pre-decision low-frequency cluster was observed around −0.9 s in the theta/alpha range, indicating that in *ideal* investors, choice-dependent neural differentiation may begin weakly prior to the decision point but becomes most pronounced after onset (38; 52). Notably, no comparably robust significant clusters emerged at the parietal channel for *ideal* investors (Fig. 3d), suggesting that their neural differentiation between decision types is expressed primarily through frontal beta-range dynamics.

*Aggressive* investors displayed the most widespread and temporally extended neural differentiation, with significant clusters present both before and after decision onset and across frontal and parietal electrodes. At the frontal channel, *aggressive* investors exhibited a prominent pre-decision cluster from approximately −1.0 to −0.5 s in the low-to-mid beta range (Fig. 3c,d), indicating anticipatory decision-specific modulation well before the decision time. Following onset, additional frontal clusters emerged in mid frequencies during the 0.1 −0.5 s window, along with a later post-decision cluster around 0.75–1.0s in beta range, reflecting sustained and multi-stage frontal engagement across the decision formation period (5; 13; 24). At the parietal channel, *aggressive* investors showed multiple significant pre-decision clusters spanning a broader frequency range, suggesting strong preparatory state differences that precede commitment. Post-decision, *aggressive* investors demonstrated a large sustained parietal cluster spanning roughly 0.1 −1.0 s in ∼10 −20 Hz. Together, these findings indicate that *aggressive* decision-making is characterized by distributed, anticipatory, and sustained oscillatory differentiation rather than a single focal post-onset signature.

*Conservative* investors showed comparatively sparse and localized effects. No significant clusters were observed at the frontal channel (Fig.3c), indicating minimal frontal differentiation between *keep* and *invest* decisions in this group. In contrast, at the parietal channel, *conservative* investors exhibited two discrete post-decision clusters in the alpha/low-beta range, occurring between about 0.2 0 −.4 s and 0.5 −0.8 s after stimulus onset (Fig.3d). This temporally constrained parietal modulation suggests that for *conservative* investors, decision differentiation is expressed primarily through posterior attentional or evidence-integration processes rather than frontal control-related dynamics, consistent with the role of parietal alpha/low-beta oscillations in controlled gating and suppression of competing representations during choice formation (16; 32).

Across strategy groups, these EEG signatures demonstrate substantial heterogeneity in the neural dynamics supporting the ambiguous decision process. *Ideal* investors show a strong and selective post-onset frontal beta-range signature, *aggressive* investors exhibit broad anticipatory and sustained differentiation across both frontal and parietal regions, and *conservative* investors show weaker effects that are localized to post-onset parietal alpha/low-beta dynamics. This pattern supports the interpretation that individual decision strategies correspond to distinct modes of neural engagement—ranging from focal frontal recruitment to distributed anticipatory oscillatory states—during strategy-dependent choice formation under ambiguity.

#### 2.4 Behavior-Physiology Coupling Reveals Divergent Internal Models of Ambiguity

When making decision under ambiguity, decision strategies diverged sharply across investor types (*F* (2, 93) = 17.66, *P <* 0.0001). *Conservative* investors exhibited a significantly higher keep-to-invest ratio than both *ideal* and *aggressive* investors (Fig. 4a), indicating a strong bias toward the safe option under moderate ambiguity. In contrast, *aggressive* investors showed the lowest keep-to-invest ratio, reflecting a pronounced tendency to commit to the risky option even when outcome probabilities were partially unknown. *Ideal* investors fell between these two extremes. This behavioral separation under matched ambiguity provides direct evidence that individual decision styles express distinct asymmetric choice policies under ambiguity.

**Fig. 4:**
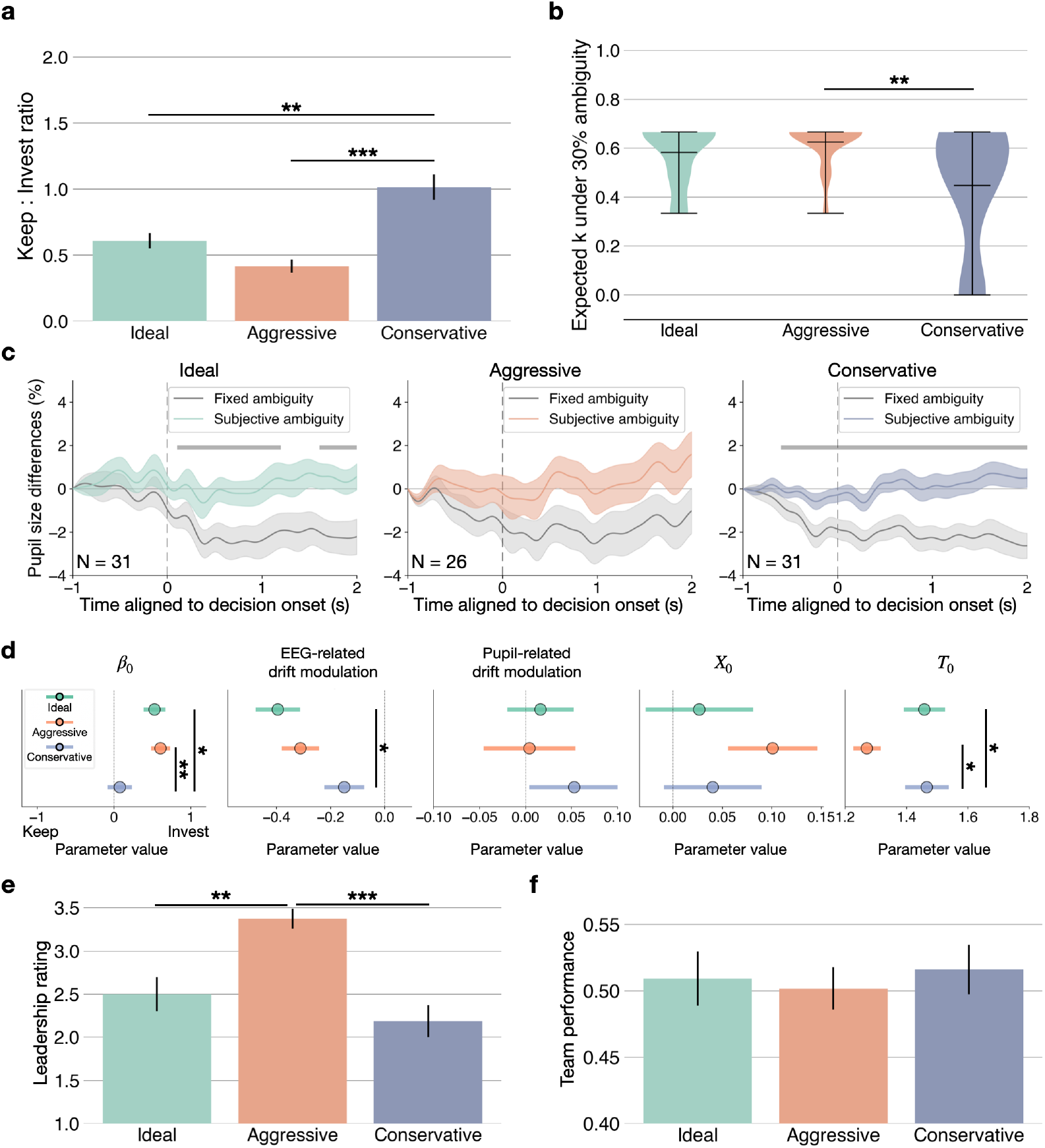
Behavioral and pupil dynamics of divergent internal models under ambiguity. **a**, Ratio of keep to invest decisions under ambiguity for *ideal, aggressive*, and *conservative* investors. *Conservative* investors exhibit a significantly higher keep-to-invest ratio than both *ideal* (*U* = 257.50, *P* = 0.0026) and *aggressive* investors (*U* = 134.00, *P <* 0.0001). **b**, Distribution of the inferred expected high-payoff probability *k* assigned to the ambiguous portion of the lottery for each investor type. *Conservative* investors assign lower values of *k* (*ideal U* = 659.50, *P* = 0.1219; *aggressive U* = 735.00, *P* = 0.0025), consistent with a pessimistic internal model of ambiguity relative to *ideal* and *aggressive* investors. Group differences in panels **a** and **b** were assessed using one-way ANOVA with post-hoc Mann-Whitney U test and Bonferroni correction (** *P <* 0.01, *\*\*\* P <* 0.001). **c**, Time-resolved pupil response difference between ambiguous and non-ambiguous trials (Ambiguity *minus* No ambiguity) for each investor type, comparing a fixed ambiguity model (equal split of high or low payoff) versus a subjective ambiguity model (each participant’s inferred high-payoff belief *Expected k*). Shaded bands indicate SEM; gray bar on top denotes significant time windows of the fixed ambiguity model (Wilcoxon signed-rank test with sliding windows and FDR correction, *P <* 0.05). **d**, Strategy-dependent drift diffusion model parameters (mean *±* SEM). Baseline drift rate (*β*_0_), EEG-related drift modulation, pupil-related drift modulation, starting point bias (*x*_0_), and non-decision time (*t*_0_) are shown for the *ideal, aggressive*, and *conservative* groups. Baseline drift differed across strategies (Kruskal-Wallis, *H*(79) = 13.46, *P* = 0.0012). EEG-related drift modulation showed a marginal group effect (*H*(79) = 5.53, *P* = 0.0629). Pupil-related drift modulation and Starting point bias (*X*_0_) did not exhibit reliable group differences. Non-decision time differed across groups (*H*(79) = 7.47, *P* = 0.0238). *N* = 25 for *ideal* and *conservative, N* = 29 for *aggressive*. **e-f**, Leadership rating and team performance from collaborative task for each characteristic of participant. Oneway Analysis of Variance (ANOVA) with post-hoc pairwise comparisons with Tukey’s HSD correction. Statistical significance is indicated as \*\**P <* 0.01, *\*\*\*P <* 0.001. N = 32 for each strategy group in a, b, d, and e.

The behavioral asymmetry was accompanied by systematic differences in how participants internally represented the ambiguous portion of the lottery (Fig. 4b). Under 30% ambiguity, the inferred expected high-payoff probability *k* differed significantly across investor types (*F* (2, 93) = 8.93, *P* = 0.0003). *Conservative* investors assigned significantly lower values of *k* than both *ideal* and *aggressive* investors, indicating that they interpreted a larger fraction of the ambiguous probability mass as yielding the low-payoff outcome (mean SEM: *ideal, k* = 0.58 *±* 0.12; *aggressive, k* = 0.63 *±* 0.09; *conservative, k* = 0.45 *±* 0.25). In contrast, *aggressive* investors assigned the highest values of *k*, reflecting a more optimistic internal model of ambiguous outcomes, with *ideal* investors again falling between the two extremes. These results demonstrate that divergent choice policies under ambiguity arise from distinct internal beliefs about the likely structure of unknown outcomes, rather than from uniform objective valuation alone.

Under a neutral valuation of ambiguity, we next examined whether these belief differences were also expressed in physiological arousal. We computed the time-resolved pupil response difference between ambiguous and non-ambiguous trials (Ambiguity *minus* No ambiguity), where ambiguity was defined using a fixed model that assumes an equal split of the ambiguous probability mass between high and low payoff outcomes. Under this neutral assumption, ambiguity elicited systematic deviations in pupil size relative to the no-ambiguity baseline (Fig. 4c, gray pupil sizes), indicating that pupil-linked arousal is modulated by ambiguity even when the objective ambiguity structure is held constant.

We then recomputed ambiguity using each participant’s inferred subjective model, parameterized by the *expected k*. Strikingly, when pupil responses were expressed in this participant-specific belief space, the ambiguity-related pupil differences were no longer significant relative to zero (*P >* 0.05; Fig.4c, colored pupil sizes). This convergence indicates that the subjective internal model inferred from behavior provides a better account of physiological arousal than a neutral equal-split rule. Together, these results support a unified interpretation in which ambiguity-driven pupil dynamics track participants’ internal beliefs about unknown outcomes: when ambiguity is quantified according to each individual’s inferred *expected k*, pupil responses become indistinguishable from the no-ambiguity baseline, consistent with the view that the behavioral and physiological signatures reflect the same latent valuation model.

#### 2.5 Divergent Evidence Accumulation Processes Drive Behavioral Strategies

To characterize latent decision dynamics underlying *keep* versus *invest* choices across the three strategies, we fit drift diffusion models (DDMs) separately for each participant in the *ideal, aggressive*, and *conservative* groups (see *Drift Diffusion Modeling of Investment Decisions* in *Materials and Methods*). Evidence accumulation evolved between a lower bound corresponding to *keep* and an upper bound corresponding to *invest*. Participant’s subjective rate of high-payoff, pupil-linked arousal, frontal–parietal EEG difference, and task context (trial progression and ambiguity) were incorporated as modulators of drift rate.

Significant strategy-dependent differences emerged in baseline drift rate (*β*_0_; Fig. 4d, *β*_0_). Post hoc Tukey HSD comparisons revealed that the *conservative* group exhibited significantly lower baseline drift relative to both the *aggressive* (*P* = 0.0036) and *ideal* (*P* = 0.020) groups, whereas the *aggressive* and *ideal* groups did not differ. These results indicate that *conservative* participants showed a reduced intrinsic tendency for evidence to accumulate toward investment, while *aggressive* and *ideal* participants exhibited comparable baseline accumulation biases toward *invest*.

Drift modulation by EEG difference showed a marginal overall group effect. Pairwise comparisons revealed weaker EEG-related drift modulation in the *conservative* group relative to the *ideal* group (*P* = 0.035). This pattern suggests that neural state fluctuations influenced evidence accumulation more strongly in *ideal* participants, whereas *conservative* participants exhibited attenuated sensitivity to EEG-derived signals. In contrast, drift modulation by pupil size did not differ significantly across strategies. This dissociation indicates that neural dynamics, rather than global arousal indexed by pupil size, primarily accounted for strategy-dependent differences in accumulation processes.

Starting point bias (*X*_0_) did not exhibit reliable group differences, indicating that pre-decisional biases were comparable across the three groups. Interestingly, Nondecision time (*t*_0_) differed significantly across groups. Tukey HSD comparisons revealed longer non-decision times in the *conservative* group relative to both the *aggressive* (*P* = 0.0306) and *ideal* (*P* = 0.0398) groups. This finding indicates that *conservative* participants required greater time for processes outside evidence accumulation than other two groups of participants.

Together, these results demonstrate that behavioral strategies are primarily distinguished by intrinsic evidence accumulation tendencies and differential neural modulation of drift. *Conservative* participants exhibited reduced baseline drift and weaker EEG-related drift modulation, consistent with diminished internal drive toward investment and attenuated neural sensitivity. *Aggressive* participants showed 14 baseline accumulation comparable to *ideal* participants but exhibited prolonged nondecision times, suggesting strategy-specific differences in peripheral processing rather than accumulation dynamics. Pupil-related drift modulation, subjective believe of high-payoff rate (*K*), and task related variables include ambiguity and trial index remained consistent across groups (Supplementary Information SFig. 2), indicating a shared computational mechanism for value integration and arousal-linked influences on decision formation.

#### 2.6 Decision Strategies and Their Correlation with Leadership and Team Performance in Collaborative Task

Having established that decision strategies correspond to stable behavioral patterns, interpretable internal beliefs, and aligned physiological signatures, we next asked whether these individual differences extend to social behavior in collaborative settings. Before the Lottery Choice Task (LCT), participants completed a three-person collaborative spacecraft control task that required real-time coordination across distinct degrees of freedom under partial and complementary visual information (48). This phase provided an independent behavioral context in which leadership tendencies and coordination styles could emerge prior to any individual ambiguity-based decisionmaking. This design enabled us to test whether the decision strategy reflects broader social traits rather than task-specific choice behavior.

Decision strategy was strongly associated with leadership behavior during collaboration. Participants classified into the three investor types received significantly different leadership ratings (*F* (2, 93) = 13.06, *p <* 0.001). *Aggressive* investors exhibited higher leadership scores than both *ideal* and *conservative* investors, whereas *conservative* investors received the lowest leadership ratings (Fig. 1b, *aggressive*-*ideal* : *P* = 0.0013; *aggressive*-*conservative*: *P <* 0.0001). This pattern suggests that individuals who assume greater leadership responsibility in team coordination are more likely to adopt risk-seeking strategies in subsequent individual value-based choices, whereas lower leadership engagement is associated with more conservative policies. These results align with prior work linking leadership, dominance, and risk tolerance in economic decision-making (20; 57).

In contrast, the decision strategy was not predictive of team performance. As shown in Fig. 1c, team performance did not significantly differ across investor types (*F* (2, 93) = 0.20, *P* = 0.820), and teams spanning the full performance range contained mixtures of *ideal, aggressive*, and *conservative* participants (Supplementary Information SFig. 1). Because performance in the collaborative task depends on the coordinated contributions of all three team members, and each potentially following a distinct decision strategy, individual investor type alone is insufficient to explain collective outcomes. This dissociation suggests that investor type captures individual leadership and risk preference, whereas team performance is dominated by interaction dynamics and coordination quality.

Together, these findings indicate that investor types reflect more than idiosyncratic ambiguity preferences: they are linked to stable social-role tendencies that generalize 15 to collaborative behavior. Leadership emerges as a key behavioral correlate of risk-seeking strategies, while team performance remains a property of the group rather than any single decision style.

## 3 Discussion

Across a value-based decision-making task with controlled ambiguity, we identified three distinct decision styles, *ideal, aggressive*, and *conservative*, that differed systematically in behavior, internal belief about ambiguous outcomes, and physiological response. *Aggressive* investors showed a faster decision-making speed, a greater tendency to select the risky option, and higher subjective expectations of receiving the high payoff. *Conservative* investors exhibited the opposite pattern: slower and fewer investment decisions, reduced sensitivity to ambiguity, and markedly lower subjective estimates of high-payoff probability. *Ideal* investors occupied an intermediate regime, showing balanced evaluation of keep and invest options and robust neural differentiation between decision types. Physiological signals, including pupil-linked arousal and EEG dynamics, further distinguished these groups and revealed that decision style is expressed not only in overt behavior but also in the underlying arousal and control processes engaged during choice. Together, these findings show that ambiguity does not evoke a uniform behavioral or physiological response across participants with different decision-making styles; instead, individuals rely on distinct internal models and computational strategies when forming decisions under ambiguity.

Classical accounts of ambiguity aversion often assume a single, population-wide tendency to avoid options with unknown probabilities (18; 14; 11). Our findings challenge this view. Individuals construct markedly different internal beliefs about ambiguous outcomes, and these beliefs shape both behavior and physiology. *Ideal, aggressive*, and *conservative* investors did not simply differ in how often they chose the risky option; they differed in their subjective estimates of the ambiguous probability mass, in their pupil-linked arousal responses, and in their neural engagement during the decision process. *Aggressive* investors assigned slightly higher subjective probability to receiving the high payoff than *ideal* investors, whereas *conservative* investors assigned much lower values, reflecting a more pessimistic internal model of ambiguity. These belief differences were mirrored in physiology, where *conservative* investors showed weaker pupil modulation by ambiguity and distinct EEG patterns of keep and invest. Taken together, these results show that ambiguity aversion is not a uniform psychological bias, but a set of heterogeneous belief-driven strategies that shape how ambiguity is represented and acted upon.

Across investor types, behavioral patterns aligned closely with participants’ internal beliefs about ambiguous outcomes (21). *Conservative* investors showed the highest keep-to-invest ratios under ambiguity and also assigned the lowest subjective k values, indicating that they not only acted cautiously but believed that high-payoff outcomes were unlikely. *Aggressive* investors displayed the opposite pattern, where they had the lowest keep-to-invest ratios and assigned the highest subjective k values. These differences suggest that *aggressive* investors held a more optimistic internal model of ambiguous outcomes than *conservative* investors. *Ideal* investors consistently fell between these two extremes. This tight correspondence between belief and behavior shows that participants’ choices were not random or inconsistent (29). Instead, they reflected stable internal valuation models that shaped how each group interpreted and acted in the face of ambiguity.

These belief–behavior links were mirrored in the physiological signatures of decision making. Under a neutral expected-value assumption, where the ambiguous probability mass was treated as evenly split between high and low pay-offs, pupil responses are significantly different across investor types when making ideal investments (Fig. 4b). This indicated that groups engaged different levels of arousal even when applying the same objective valuation rule (7; 60; 8). However, these differences disappeared once the expected value was recomputed using each participant’s own subjective *k* (Fig. 4c). When ideal decisions were evaluated under each individual’s internal belief model, pupil responses no longer differed across groups. This shift demonstrates that pupil-linked arousal does not reflect a fixed reaction to ambiguity across groups. Instead, it tracks the subjective value each participant assigns to the ambiguous portion, aligning directly with their internal beliefs and behavioral tendencies. Physiological responses, therefore, provide convergent evidence that ambiguity processing is belief-driven, not solely stimulus-driven.

Neural dynamics further revealed that each investment style engages a distinct computational pathway when resolving ambiguity. *Ideal* investors showed early enhancements in delta and theta power, along with broader frontoparietal recruitment, consistent with rapid evaluation of competing options and early engagement of control systems (61; 43; 55). In contrast, *aggressive* and *conservative* investors showed limited early low-frequency activity and instead differentiated keep and invest decisions through posterior alpha modulation, reflecting shifts in cortical excitability and attentional allocation (51; 59). *Aggressive* investors additionally showed a late alpha signature when choosing against their default tendency, suggesting engagement of conflict-monitoring or effortful control processes (41; 42). *Conservative* investors, by comparison, displayed generally muted low-frequency responses, consistent with a less dynamically engaged decision regime. Together, these patterns indicate that individuals do not simply vary in their decision thresholds; they draw on qualitatively different neural mechanisms when forming decisions under ambiguity.

Across these modalities, a unified model of ambiguity processing emerges. *Ideal* investors combine balanced subjective beliefs with early control-related neural engagement, enabling them to evaluate ambiguous options through an integrated valuation process. *Aggressive* investors display the opposite pattern: they hold more optimistic beliefs about ambiguous outcomes and rely on heightened cortical excitability to commit rapidly to the risky option, engaging additional control signals only when acting against their default tendency (41; 42). *Conservative* investors adopt a pessimistic interpretation of ambiguity and operate in a low-engagement neural regime, with weaker arousal responses and muted low-frequency signaling that track their reluctance to invest. Together, these profiles suggest that individuals do not differ merely in their preferences under ambiguity. Different groups differ in the internal models they construct, the arousal systems they recruit, and the neural computations they engage when forming decisions. Ambiguity processing, therefore, reflects distinct belief-driven pathways rather than a single canonical mechanism.

Our findings also refine several longstanding theoretical accounts of decision making under ambiguity. Classical ambiguity frameworks, beginning with Ellsberg (18), assume a uniform population-level aversion to unknown probabilities, but our results show that this aggregate pattern masks structured heterogeneity in how individuals internally represent ambiguity (Fig. 2a). Our data demonstrate that internal beliefs about receiving high payoffs are reflected not only in choice patterns but also in arousal signatures and neural dynamics. Work on LC–NE–mediated arousal has highlighted pupil dilation as a marker of ambiguity or cognitive demand (62; 45; 23), but our results show that pupil responses also track subjective valuation rather than just ambiguity. In decision neuroscience, theta, alpha, and delta rhythms are often interpreted as generic markers of control, attention, or valuation (25; 63; 10; 61). We show that participants’ expressions differ systematically across decision styles, revealing distinct computational modes rather than simple increases or decreases in activation. Together, these links to prior theory demonstrate that ambiguity is not a single psychological construct but a multi-level phenomenon shaped by individual beliefs, physiological engagement, and neural computation.

Importantly, our results also indicate that decision-making under ambiguity is shaped not only by stimulus-evoked valuation processes but also by pre-stimulus internal states. In pupillometry, all three strategy groups exhibited reliable *keep* - *invest* differences prior to stimulus onset, suggesting that endogenous arousal fluctuations can predict forthcoming choice (56; 17). EEG provided convergent, but strategy-dependent, evidence for such anticipatory differentiation. *Aggressive* investors showed the clearest pre-stimuli differences between *keep* versus *invest* decisions, indicating that neural states predictive of choice can emerge well before the decision period (38; 52). *Ideal* investors exhibited only a weaker pre-decision effect, and *conservative* investors showed no significant pre-decision EEG clusters. These results suggest that anticipatory neural differentiation is not uniformly expressed across participants with strategies. One possibility is that participants enter each trial with fluctuating internal states, such as expectation, decision bias, or action preparedness, that bias the probability of selecting *keep* versus *invest*. Alternatively, these pre-stimulus physiological differences may reflect carryover from the previous trial, producing lingering engagement or subjective certainty that biases the subsequent decision. Together, these findings suggest that ambiguity processing unfolds within a dynamically varying internal context, in which endogenous arousal and preparatory neural states shape decision outcomes even before evidence is evaluated.

Several limitations of this study also point to clear directions for future work. First, our laboratory task used fixed ambiguity levels (30% and 60%), which enabled precise inference of internal belief models but may not capture the full complexity of real-world ambiguity. Extending this framework to naturalistic settings, such as real-time financial decision-making, including stock trading or shopping decisions, will be an important next step. Second, although pupil size reliably indexed arousal and tracked subjective valuation, it is not a process-pure marker of LC-NE activity. Future studies combining pupillometry with higher-resolution methods, such as EEG–fMRI, could more directly link arousal dynamics to neuromodulatory mechanisms. Third, the limited spatial resolution of EEG constrains source-level interpretation, motivating multimodal approaches to better resolve the neural circuits underlying belief-driven ambiguity processing. More broadly, our results open avenues for linking internal belief models to personality, risk preference, and psychiatric phenotypes, and for designing adaptive decision-support and human–AI systems that respond to latent beliefs rather than observable choices alone.

## 4 Materials and Methods

### 4.1 Participants

We recruited 57 healthy adult participants (mean age 23.68 *±* 3.32 years; 25 female), all with normal or corrected-to-normal vision. Each individual completed one to three experimental sessions. For data analysis, each session was treated as an independent participant, yielding a total of 108 sessions. Data from 7 sessions were excluded due to incompletion, and 5 additional sessions were excluded due to low task engagement (selecting only one option across trials). EEG data from 17 sessions were discarded due to poor signal quality, and pupil size data from 1 sessions were excluded due to failed eye-tracker calibration. The final dataset included behavioral data from 96 sessions, pupil data from 95 sessions, and EEG data from 79 sessions. All procedures were approved by the Institutional Review Board at Columbia University, and written informed consent was obtained from all participants prior to each session.

### 4.2 Experimental Design

Each participant completed the Lottery Choice Task (LCT) following a collaborative decision-making task, the Apollo Distributed Control Tast, and an associated questionnaire (46; 47; 48). The LCT was designed to assess individual risk preferences under varying levels of ambiguity and under time pressure. Participants were instructed to maximize their total score, which was partially linked to monetary rewards.

Each trial began with a 1 second fixation period, followed by a decision phase lasting up to 5 seconds. Participants were randomly assigned an initial point endowment and asked to either keep the amount or invest it for a probabilistic return. The repayment probability was displayed using a pie chart representing either (1) a social condition, featuring a human face labeled as a prior participant, or (2) a non-social condition, featuring a roulette image representing a computer-generated gamble. There was no correlation between stimulus type and the probability of return.

After the decision was made and confirmed, a 2 second reviewing period began. During this phase, a ball rotated around the outer pie chart and stopped to reveal the investment outcome, indicated both by the ball’s final color and an outcome message displayed above the pie chart. If participants failed to respond in time, the trial was flagged as skipped and repeated later.

The task manipulated three levels of ambiguity:

- **No ambiguity:**all outcome probabilities and amounts were fully known.
- **Moderate ambiguity:** 30% of the pie chart was hidden.
- **High ambiguity:** 60% of the pie chart was hidden.

In ambiguous trials, the hidden region could belong entirely or partially to either payoff region, creating uncertainty about both the likelihood and the value of the outcome.

Each session contained 132 valid trials. Trials in which participants failed to respond within the 5 s decision window were repeated later at a random position in the session. Therefore, the total number of trials presented could exceed 132, but 132 trials were retained for analysis. The task was implemented in an immersive virtual reality environment. Participants interacted with the task using a head-mounted display (HTC VIVE Pro Eye, 1440 *×* 1600 pixels per eye, refresh rate of 90 Hz) and handheld controllers (HTC VIVE Controller) while seated in a quiet, EEG-shielded chamber. Behavioral responses, eye tracking (pupil size), and EEG signals were recorded continuously throughout the task.

The virtual interface displayed two side-by-side panels per trial. One panel showed a full pie displaying the initial endowment. Another panel displayed the stimulus (face or roulette) above a pie chart indicating repayment probabilities and amounts. Each segment of the pie showed a numerical value representing the potential return and a corresponding percentage around the outer edge. In ambiguous conditions, a portion of the pie was shaded without a numerical label, and only the percentage of the unknown portion was displayed. Participants confirmed their selection by clicking a “submit” button located between the two panels after each trial. The reviewing period began immediately after submission.

### 4.3 Behavioral Data and Participants Classification

To understand how participants made investment decisions under varying conditions, we analyzed the frequency of “keep” versus “invest” responses across different levels of ambiguity and stimulus type. We then used physiological signals, combined with game context variables, to build predictive models of individual investment decisions.

Each investment trial was associated with an expected value (EV), computed as:

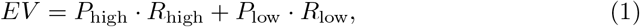

where *P* denotes the probability and *R* the corresponding return amount. For ambiguous trials, EV was evaluated using the actual underlying probabilities excluding the ambiguous portion.

We categorized each decision into one of three types based on the comparison between the EV and the initial endowment:

- **Ideal decision:** Participant chose to invest when *EV >* initial, or to keep when *EV <* initial.
- **Aggressive decision:** Participant chose to invest even when *EV <* initial.
- **Conservative decision:** Participant chose to keep even when *EV >* initial.

For each participant, we computed the proportion of decisions falling into each category. Participants were then ranked by the proportion of ideal, aggressive, and conservative decisions. The top one-third were labeled as *Ideal investors*. Among the remaining two-thirds, the top half (one-third overall) were labeled as *Conservative investors*, based on their conservative decision rate. The remaining participants were categorized as *Aggressive investors*.

This classification yielded three mutually exclusive behavioral profiles, which served as the foundation for subsequent physiological and behavioral analyses and predictive modeling.

### 4.4 Pupil Data Acquisition and Preprocessing

Pupil data were recorded using the integrated eye tracker on the HTC VIVE Pro Eye virtual reality headset, operating at a sampling rate of 120 Hz with a trackable field of view of 110^◦^. The system recorded pupil size, gaze position, gaze direction, and eye openness continuously throughout the task.

Pupil size preprocessing involved blink and artifact detection, followed by linear interpolation over missing values. The interpolated signal was then filtered using a 4th-order Butterworth low-pass filter with a cutoff frequency of 2 Hz. Pupil diameter was averaged across the two eyes to obtain a single time series per trial.

Pupil size was normalized for each trial using the formula:

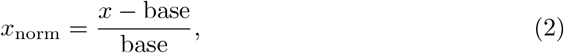

where *x* is the pupil size time series and base is the baseline pupil size measured during the 1-second fixation period preceding the decision phase. We analyzed pupil dynamics during the 0–2 second window following stimulus onset, corresponding to the decision-making period. Trials with decision times shorter than 2 seconds were excluded from pupil analysis to ensure consistent temporal alignment across trials.

To isolate decision-related pupil dynamics, we removed the transient pupil response evoked by stimulus luminance changes using a generalized linear model (GLM). For each participant, we modeled the luminance-evoked pupil response as a step-like impulse response using a finite impulse response (FIR) basis and fit the model only to the early post-stimulus window (the first 1 second), when luminance-driven effects dominate. A ridge-regularized regression was used to estimate a participant-specific luminance response kernel shared across trials. The fitted luminance component was then extrapolated to the full trial, held constant after the early window to avoid removing later decision-related dynamics, and smoothly blended at the boundary. This estimated luminance contribution was subtracted from the raw pupil signal, yielding a luminance-corrected pupil trace used for all subsequent analyses.

### 4.5 EEG Data Acquisition and Analysis

EEG data were recorded using the B-Alert X24 system (Advanced Brain Monitoring), which includes 20 electrodes placed according to the international 10–20 system and sampled at 256 Hz. The system used two mastoid electrodes for referencing. Participants completed the task while seated in an EEG-shielded booth to minimize environmental artifacts.

EEG preprocessing was performed using custom Python scripts. First, bad channels were detected and excluded based on signal amplitude thresholds using PyPREP (2). The removed channels are interpolated using MNE (28). The data were then bandpass-filtered between 0.5 and 30 Hz using a zero-phase Butterworth filter. Independent component analysis (ICA) was applied to identify and remove components related to ocular and muscular artifacts. All EEG traces were visually inspected post-ICA to confirm artifact removal.

Continuous EEG data were epoched around stimulus onset, spanning a window from 0.25 to 1 seconds. The −0.25 *s* preceding stimulus onset served as the baseline for baseline correction and was excluded from subsequent analyses. All analyses focused on the post-stimulus interval from 0 to 1 second. Participants with fewer than 5 valid trials per condition were excluded from EEG analysis. Following baseline correction using the pre-stimulus window (−0.25 to 0 s), we cropped epochs to the 0–1 second post-stimulus interval.

Frequency-domain analysis targeted the delta (0.5–4 Hz), theta (4–8 Hz), and alpha (8–13 Hz) bands, which are strongly implicated in cognitive control and value-based decision making (39; 53).

### 4.6 Drift Diffusion Modeling of Investment Decisions

To quantify latent cognitive processes underlying *keep* and *invest* decisions (49), we employed drift diffusion modeling (DDM) using the PyDDM framework (54). Choices were coded as keep = 0 and invest = 1, corresponding to the lower and upper absorbing decision bounds, respectively. Models were fit separately for each participant using trial-wise decision times and binary choices.

Trial-wise EEG features were computed using Morlet wavelet time-frequency decomposition implemented in MNE (28). Frequencies from 0.5 to 30 Hz (60 linearly spaced bins) were used with the number of cycles set to half the frequency. Power estimates were baseline-normalized using a log-ratio transform relative to a pre-stimulus baseline window of [−0.7, −0.5] s. For each trial, baseline-normalized power was averaged within the 8-13 Hz band and a 0-0.5 s post stimuli onset time window at channels Fz and Pz. The EEG regressor used in the model was defined as the difference between these two channels,

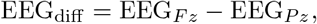

capturing trial-wise frontal-parietal spectral imbalance.

The subjective value term *K* was assigned according to each participant’s internal believe of the high-payoff rate under ambiguity, yielding a participant-specific estimate of expected value under uncertainty. Trial-wise pupil and EEG covariates were z-score normalized within each participant to remove between-subject scaling differences.

Evidence accumulation was modeled as a stochastic diffusion process evolving between two fixed absorbing bounds. The diffusion noise was fixed to 1.0 and the boundary separation was fixed to 1.8 across participants. The model estimated a starting point parameter (*x*_0_), representing initial decision bias toward either choice, and a non-decision time parameter (*t*_0_), capturing sensory encoding and motor execution processes outside the accumulation stage.

The drift rate was modeled as a linear function of trial-level cognitive and physiological covariates:

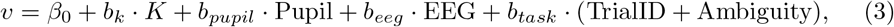

where *β*_0_ denotes the baseline drift rate, *K* denotes subjective value of getting high-payoff, Pupil denotes the trial-wise pupil scalar, EEG denotes the frontal-parietal 13-15 Hz EEG difference in 0.1 to 0.4 s post stimuli onset, and TrialID + Ambiguity encodes the trial index and ambiguity level of the trial. Positive drift values indicate faster accumulation toward the *invest* decision.

Model parameters were estimated independently for each participant using maximum-likelihood estimation with the BADS optimizer implemented in PyDDM. Parameter bounds were constrained to ensure stable optimization: *β*_0_, *b*_*k*_, *b*_*pupil*_, *b*_*eeg*_, *b*_*task*_, [−1, 1], *x*_0_ ∈ [−1, 1], bound ∈ [0, 2], and *t*_0_ ∈ [0, 2]. This procedure yielded participant-specific estimates of baseline accumulation tendencies, sensitivity to subjective value, modulation by pupil-linked arousal, EEG-derived neural state, task related effects, starting bias, bound, and non-decision time. The model fit quality measured by negative log likelihood is shown in Supplementary Information SFig. 3.

### 4.7 Statistical Analyses

#### 4.7.1 Behavioral

To verify that differences in investment decision patterns across ambiguity levels and stimulus types are statistically significant, we used Cochran–Mantel–Haenszel (CMH) tests to evaluate stratified contingency tables, controlling for participant-level variation. Reaction times were analyzed using a linear mixed-effects model with stimulus type as a fixed effect and a random intercept for each participant:

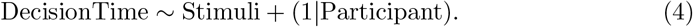

For group-level comparisons related to leadership scores and team performance (derived from the preceding Apollo Distributed Control Task), we applied one-way ANOVA with Bonferroni correction for multiple comparisons.

#### 4.7.2 Physiological

Pupil size time-resolved condition effects were evaluated using sliding-window tests across the pupil time course (0.4 s window; 0.2 s step). For within-participant comparisons (e.g., *keep* vs. *invest* within an investor type), we used Wilcoxon signed-rank tests on the window-averaged pupil values. For between-group comparisons across investor types (*ideal, aggressive*, and *conservative*) at each window, we used Kruskal–Wallis tests. For each analysis, we controlled multiple comparisons across windows using FDR correction, and significant time intervals were reported/visualized as contiguous FDR-significant windows.

Statistical comparisons between *keep*and *invest* decisions’ EEG were performed using paired tests across participants in a sliding-window manner over time and frequency. Specifically, power values were averaged within overlapping temporal windows (0.1 s windows with 0.05 s steps) and narrow frequency bands (3 Hz width). Paired-sample tests were computed for each time–frequency bin, and false discovery rate (FDR) correction was applied across all bins to control for multiple comparisons.

## Acknowledgements

This work was supported by funding from the Army Research Laboratory’s STRONG Program (W911NF-19-2-0139, W911NF-19-2-0135, W911NF-21-2-0125), the National Science Foundation (IIS-1816363, OIA-1934968), the Air Force Office of Scientific Research (FA9550-22-1-0337), and a Vannevar Bush Faculty Fellowship from the US Department of Defense (N00014-20-1-2027).

## References

[1] Al-Najjar NI, Weinstein J (2009) The ambiguity aversion literature: a critical assessment. Economics & Philosophy 25(3):249–284

[2] Bigdely-Shamlo N, Mullen T, Kothe C, et al (2015) The prep pipeline: standardized preprocessing for large-scale eeg analysis. Frontiers in neuroinformatics 9:16

[3] Blankenstein NE, Peper JS, Crone EA, et al (2017) Neural mechanisms underlying risk and ambiguity attitudes. Journal of Cognitive Neuroscience 29(11):1845–1859

[4] Blankenstein NE, Huettel SA, Li R (2021) Resolving ambiguity: Broadening the consideration of risky decision making over adolescent development. Developmental Review 62:100987

[5] Bode S, Sewell DK, Lilburn S, et al (2012) Predicting perceptual decision biases from early brain activity. Journal of Neuroscience 32(36):12488–12498

[6] Botelho C, Fernandes C, Campos C, et al (2023) Uncertainty deconstructed: conceptual analysis and state-of-the-art review of the erp correlates of risk and ambiguity in decision-making. Cognitive, Affective, & Behavioral Neuroscience 23(3):522–542

[7] Bradshaw J (1967) Pupil size as a measure of arousal during information processing. Nature 216(5114):515–516

[8] Carro-Domínguez M, Huwiler S, Oberlin S, et al (2025) Pupil size reveals arousal level fluctuations in human sleep. Nature communications 16(1):2070

[9] Cavanagh JF, Frank MJ (2014) Frontal theta as a mechanism for cognitive control. Trends in cognitive sciences 18(8):414–421

[10] Cavanagh JF, Shackman AJ (2015) Frontal midline theta reflects anxiety and cognitive control: meta-analytic evidence. Journal of physiology-Paris 109(1-3):3–15

[11] Cubitt R, Van De Kuilen G, Mukerji S (2020) Discriminating between models of ambiguity attitude: A qualitative test. Journal of the European Economic Association 18(2):708–749

[12] De Gee JW, Knapen T, Donner TH (2014) Decision-related pupil dilation reflects upcoming choice and individual bias. Proceedings of the National Academy of Sciences 111(5):E618–E625

[13] Devine CA, Gaffney C, Loughnane GM, et al (2019) The role of premature evidence accumulation in making difficult perceptual decisions under temporal uncertainty. Elife 8:e48526

[14] Dimmock SG, Kouwenberg R, Mitchell OS, et al (2016) Ambiguity aversion and household portfolio choice puzzles: Empirical evidence. Journal of Financial Economics 119(3):559–577

[15] Einhorn HJ, Hogarth RM (1986) Decision making under ambiguity. Journal of business pp S225–S250

[16] Eldar E, Cohen JD, Niv Y (2013) The effects of neural gain on attention and learning. Nature neuroscience 16(8):1146–1153

[17] Eldar E, Felso V, Cohen JD, et al (2021) A pupillary index of susceptibility to decision biases. Nature Human Behaviour 5(5):653–662

[18] Ellsberg D (1961) The crude analysis of strategy choices. The American Economic Review 51(2):472–478

[19] Epishin V, Bogacheva N (2020) Tolerance for uncertainty and patterns of decision-making in complex problem-solving strategies. Multimodal Technologies and Interaction 4(3):58

[20] Ertac S, Gurdal MY (2012) Deciding to decide: Gender, leadership and risk-taking in groups. Journal of Economic Behavior & Organization 83(1):24–30

[21] Fairley K, Vyrastekova J, Weitzel U, et al (2019) Subjective beliefs about trust and reciprocity activate an expected reward signal in the ventral striatum. Frontiers in Neuroscience 13:660

[22] FeldmanHall O, Glimcher P, Baker AL, et al (2016) Emotion and decision-making under uncertainty: Physiological arousal predicts increased gambling during ambiguity but not risk. Journal of Experimental Psychology: General 145(10):1255

[23] Filipowicz AL, Glaze CM, Kable JW, et al (2020) Pupil diameter encodes the idiosyncratic, cognitive complexity of belief updating. Elife 9:e57872

[24] Forster C, Stephani T, Grund M, et al (2025) Pre-stimulus beta power mediates explicit and implicit perceptual biases in distinct cortical areas. Communications Psychology 3(1):93

[25] Foxe JJ, Snyder AC (2011) The role of alpha-band brain oscillations as a sensory suppression mechanism during selective attention. Frontiers in psychology 2:154

[26] Geng JJ, Blumenfeld Z, Tyson TL, et al (2015) Pupil diameter reflects uncertainty in attentional selection during visual search. Frontiers in human neuroscience 9:435

[27] Gluth S, Rieskamp J, Büchel C (2013) Classic eeg motor potentials track the emergence of value-based decisions. Neuroimage 79:394–403

[28] Gramfort A, Luessi M, Larson E, et al (2013) MEG and EEG data analysis with MNE-Python. Frontiers in Neuroscience 7(267):1–13. 10.3389/fnins.2013.00267

[29] Granados Samayoa JA, Albarracín D (2025) Understanding belief-behavior correspondence: Beliefs and belief-to-behavior inferences. Psychological Inquiry 36(1):1–22

[30] Graves JE, Egré P, Pressnitzer D, et al (2021) An implicit representation of stimulus ambiguity in pupil size. Proceedings of the National Academy of Sciences 118(48):e2107997118

[31] Guo M, Ikink I, Roelofs K, et al (2025) Ambiguity preferences in intertemporal and risky choice: A large-scale study using drift-diffusion modelling. Psychonomic Bulletin & Review pp 1–18

[32] Klimesch W (2012) Alpha-band oscillations, attention, and controlled access to stored information. Trends in cognitive sciences 16(12):606–617

[33] Kocher MG, Lahno AM, Trautmann ST (2018) Ambiguity aversion is not universal. European Economic Review 101:268–283

[34] Krain AL, Wilson AM, Arbuckle R, et al (2006) Distinct neural mechanisms of risk and ambiguity: a meta-analysis of decision-making. Neuroimage 32(1):477–484

[35] Larsen T, O’Doherty JP (2014) Uncovering the spatio-temporal dynamics of value-based decision-making in the human brain: a combined fmri–eeg study. Philosophical Transactions of the Royal Society B: Biological Sciences 369(1655):20130473

[36] Lauharatanahirun N, Aimone JA, Gately JB (2025) Risk behind the veil of ambiguity: Decision-making under social and nonsocial sources of uncertainty. Risk Analysis 45(10):3144–3159

[37] Lauriola M, Levin IP, Hart SS (2007) Common and distinct factors in decision making under ambiguity and risk: A psychometric study of individual differences. Organizational Behavior and Human Decision Processes 104(2):130–149

[38] Lou B, Hsu WY, Sajda P (2015) Perceptual salience and reward both influence feedback-related neural activity arising from choice. Journal of Neuroscience 35(38):13064–13075

[39] Nácher V, Ledberg A, Deco G, et al (2013) Coherent delta-band oscillations between cortical areas correlate with decision making. Proceedings of the National Academy of Sciences 110(37):15085–15090

[40] O’connell RG, Dockree PM, Kelly SP (2012) A supramodal accumulation-to-bound signal that determines perceptual decisions in humans. Nature Neuroscience 15(12):1729–1735

[41] Pastötter B, Hanslmayr S, Bäuml KHT (2010) Conflict processing in the anterior cingulate cortex constrains response priming. NeuroImage 50(4):1599–1605

[42] Pastötter B, Dreisbach G, Bäuml KHT (2013) Dynamic adjustments of cognitive control: oscillatory correlates of the conflict adaptation effect. Journal of Cognitive Neuroscience 25(12):2167–2178

[43] Pierrieau E, Kessouri S, Lepage JF, et al (2022) Theta but not beta activity is modulated by freedom of choice during action selection. Scientific Reports 12(1):9115

[44] Pisauro MA, Fouragnan E, Retzler C, et al (2017) Neural correlates of evidence accumulation during value-based decisions revealed via simultaneous eeg-fmri. Nature communications 8(1):15808

[45] Preuschoff K, ‘t Hart BM, Einhäuser W (2011) Pupil dilation signals surprise: Evidence for noradrenaline’s role in decision making. Frontiers in neuroscience 5:115

[46] Qin Y, Zhang W, Lee R, et al (2022) Predictive power of pupil dynamics in a team based virtual reality task. In: 2022 IEEE Conference on Virtual Reality and 3D User Interfaces Abstracts and Workshops (VRW), IEEE, pp 592–593

[47] Qin Y, Zhang W, Lee R, et al (2024) Pupil-linked arousal is predictive of team performance in a virtual reality (vr) sensory-motor task. In: 2024 46th Annual International Conference of the IEEE Engineering in Medicine and Biology Society (EMBC), IEEE, pp 1–4

[48] Qin Y, Lee RT, Zhang W, et al (2025) Physiologically informed predictability of a teammate’s future actions forecasts team performance. iScience 28(5)

[49] Ratcliff R, McKoon G (2008) The diffusion decision model: theory and data for two-choice decision tasks. Neural computation 20(4):873–922

[50] Rungratsameetaweemana N, Itthipuripat S, Salazar A, et al (2018) Expectations do not alter early sensory processing during perceptual decision-making. Journal of Neuroscience 38(24):5632–5648

[51] Sauseng P, Feldheim JF, Freunberger R, et al (2011) Right prefrontal tms disrupts interregional anticipatory eeg alpha activity during shifting of visuospatial attention. Frontiers in psychology 2:241

[52] Schmidt R, Ruiz MH, Kilavik BE, et al (2019) Beta oscillations in working memory, executive control of movement and thought, and sensorimotor function. Journal of Neuroscience 39(42):8231–8238

[53] Senftleben U, Scherbaum S (2021) Mid-frontal theta during conflict in a value-based decision task. Journal of Cognitive Neuroscience 33(10):2109–2131

[54] Shinn M, Lam NH, Murray JD (2020) A flexible framework for simulating and fitting generalized drift-diffusion models. ELife 9:e56938

[55] Thornberry C, Commins S (2024) Frontal delta and theta power reflect strategy changes during human spatial memory retrieval in a virtual water maze task: an exploratory analysis. Frontiers in Cognition 3:1393202

[56] Urai AE, Braun A, Donner TH (2017) Pupil-linked arousal is driven by decision uncertainty and alters serial choice bias. Nature communications 8(1):14637

[57] Van Kleef GA, Heerdink MW, Cheshin A, et al (2021) No guts, no glory? how risk-taking shapes dominance, prestige, and leadership endorsement. Journal of applied psychology 106(11):1673

[58] Viglione A, Mazziotti R, Pizzorusso T (2023) From pupil to the brain: New insights for studying cortical plasticity through pupillometry. Frontiers in neural circuits 17:1151847

[59] Wang C, Rajagovindan R, Han SM, et al (2016) Top-down control of visual alpha oscillations: sources of control signals and their mechanisms of action. Frontiers in human neuroscience 10:15

[60] Wang CA, Baird T, Huang J, et al (2018) Arousal effects on pupil size, heart rate, and skin conductance in an emotional face task. Frontiers in neurology 9:1029

[61] Wang J, Chen Z, Peng X, et al (2016) To know or not to know? theta and delta reflect complementary information about an advanced cue before feedback 28 in decision-making. Frontiers in psychology 7:1556

[62] Van der Wel P, Van Steenbergen H (2018) Pupil dilation as an index of effort in cognitive control tasks: A review. Psychonomic bulletin & review 25(6):2005–2015

[63] Wyart V, De Gardelle V, Scholl J, et al (2012) Rhythmic fluctuations in evidence accumulation during decision making in the human brain. Neuron 76(4):847–858

[64] Zhang S, Tian Y, Liu Q, et al (2024) The neural correlates of ambiguity and risk in human decision-making under an active inference framework. eLife

